# Estimating Daily Taxon-specific Tree Pollen at a 1-km Resolution in Atlanta, GA from 2020 to 2024

**DOI:** 10.64898/2026.05.14.725192

**Authors:** Xueying Zhang, Wenhao Wang, Yohei Saburi, Hannah R. Paduch, Zhihao Jin, Kai Zhu, Yang Liu

**Author notes:** Corresponding author: Xueying Zhang, Center for Precision Environmental Health, Baylor College of Medicine, Houston, TX, USA.

## Abstract

While tree pollen is a major trigger of allergic respiratory conditions and different taxa exhibit varying allergenic potentials, the lack of high-resolution, taxon-specific exposure metrics have limited our ability to identify which local pollen taxa are primarily responsible for respiratory illness. Traditional pollen monitoring networks, which have an intermittent sampling schedule, are not ideal for examining the delayed effects of pollen exposure due to the missing days. In this study, we developed a modeling framework integrating atmospheric dispersion effects, taxa-specific phenology, and machine learning to predict daily counts of 13 tree taxa in the five-county Metro Atlanta area, Georgia at a 1-km resolution from 2020 to 2024. Machine learning model performance was validated with daily pollen counts collected by a multi-site monitoring network equipped with automated pollen sensors. Findings showed that *Betula* and *Quercus* pollens exhibited higher predictive performance than other taxa, with R^2^ values ranging from 0.69 to 0.92 and from 0.71 to 0.89, respectively. Our 1-kilometer prediction data provides gapless exposure metrics to understand the spatial and temporal variability in pollen exposure, can facilitate investigation of urban pollen hotspots and support epidemiologic studies of pollen-related respiratory outcomes.

## Introduction

Tree pollen is a major trigger of allergic respiratory conditions and can cause more severe health outcomes in vulnerable populations, such as young children or older adults with existing chronic respiratory conditions.^1-3^ The species responsible for respiratory illness are suspected to vary substantially across locations due to the spatial heterogeneity in tree distributions and their particular allergenic potentials. This has been observed in epidemiologic studies examining pediatric asthma exacerbations with multiple tree pollen taxa across geographic locations.^4-7^ Among those studies, sixteen tree taxa were evaluated. The taxa associated with asthma outcomes differed by study, likely due to the different study regions; notably, *Quercus* (oak), a tree widely used in urban planning, was the only taxon consistently associated with pediatric asthma exacerbations.^4-7^ Moreover, sensitization studies from Europe,^8, 9^ Asia,^10, 11^ and the Asia-Pacific^12^ further confirmed geographic heterogeneity in pollen exposures. Despite this, most epidemiologic studies still rely on single-site pollen measurements and lack high-resolution exposure data to capture locally dominant pollen species.

Pollen exposure assessments have traditionally relied on observational networks. The most widely used pollen data source in the U.S. is the National Allergy Bureau (NAB), operated by the American Academy of Allergy, Asthma, and Immunology. To date, the NAB only includes about 50 active sites across the continental U.S., typically one per major metropolitan area. Sites in this network employ Rotorod or Burkard samplers to collect airborne particles on filters, which are then manually identified and counted under a microscope. Such an approach is highly labor-intensive and, as a result, most NAB sites do not collect samples on weekends or holidays. NAB’s discontinuous sampling schedule makes it difficult for epidemiologic studies to accurately examine the respiratory impacts of pollen exposures, as hospital visits may resulting from pollen allergens occur one to two days after exposure.^7, 13, 14^

Geospatial modeling offers a practical solution for estimating airborne pollutants at times and locations not covered by ground monitors. A few research groups have developed geospatial models for taxon-specific pollen estimates, using statistical methods,^15-20^ machine learning,^21-23^ and chemical transport models.^24-28^ The development of these geospatial models has primarily focused on two key components: the phenological patterns of pollen release and the atmospheric dispersion processes. Phenology describes the temporal variation of pollen during pollen season, including the season start date and length, peak pollen date, and the amount of pollen emitted each day, which has been extensively studied in plant ecology. Atmospheric dispersion processes estimate pollen trajectory from release to deposition under different meteorological conditions and landscapes. In recent years, phenological models have increasingly been integrated with atmospheric dispersion models to jointly capture the spatial and temporal variability of airborne pollen.

We summarize relevant studies that have advanced this integrated modeling paradigm. Zhang et al.^28^ used spatially resolved surface temperatures from the Weather Research and Forecasting (WRF) model to estimate the season start dates and lengths of five allergenic pollens (birch, oak, ragweed, mugwort, and grass) in the U.S. at a 50-m spatial resolution. The same group^26, 27^ then applied the Community Multiscale Air Quality (CMAQ) modeling system and the Hybrid Single-Particle Lagrangian Integrated Trajectory model (HYSPLIT) to predict taxon-specific pollen at 12-km^27^ and 36-km^25^ spatial resolutions, accounting for atmospheric dispersion trajectories and meteorological interactions.

Lo et al.^29^ and Song et al.^30^ developed fine-scale phenology indexes using Enhanced Vegetation Index (EVI) data retrieved from the Moderate Resolution Imaging Spectroradiometer (MODIS) satellite and the PlanetScope satellite, respectively. In Lo et al., ^29^ temporal variations in MODIS EVI, estimated green-up dates, and other climate-related variables (e.g., temperature) were used as inputs in a random forest model to predict pollen concentrations across various timespans. The model’s inclusion of prior-day pollen measurements as a predictor led to that predictions are only available in locations with existing pollen monitors. Notably, models that did not include prior-day pollen showed poor performance in their analyses.

Song et al.^30^ extended Lo et al.’s work by fitting phenological and climatic parameters directly to pollen observations, generating a time series proxy for daily pollen emission trends. By leveraging the high spatial resolution of PlanetScope images, the generated phenological time series can reach a three-meter spatial resolution, which enables pollen variability prediction at the level of individual trees. Additionally, Wozniak and Steiner^24^ developed a Pollen Emissions for Climate Models (PECM) modeling system integrating geography, vegetation type, and meteorological variables and produced climate-sensitive phenology estimates for multiple pollen taxa. PECM can simulate pollen emissions at a varied time scale and 25-km spatial resolution with nationwide coverage. Their estimated pollen emission factors have been applied in several epidemiological studies to investigate taxon-specific associations with respiratory health outcomes.^1, 31, 32^ Together, these studies have advanced pollen modeling by integrating methods in phenology, emissions, and large-scale transport, and by leveraging remote sensing and chemical transport models across a range of spatial resolutions. However, key limitations include the reliance on observed pollen measurements, which restricts pollen predictions in areas without monitoring data, and coarse spatial resolution that limits the ability to capture local variation in taxa.

In this study, we aimed to develop a scalable modeling framework to predict daily exposure estimates for 13 tree pollen taxa across the five-county Atlanta metropolitan area. Atlanta provides an ideal study setting due to its high tree density and the presence of an established pollen monitoring network that provides validation data. Details on the development of model predictors and the validation datasets are described in the following sections.

## Materials and Methods

We developed a hybrid modeling framework (**Figure 1**) to estimate taxon-specific pollen concentrations by integrating automated sensor data, geospatial modeling, and machine learning approaches. The resulting exposure estimates were generated at a 1-km spatial resolution, enabling characterization of neighborhood-level variability. Outputs from the Gaussian plume dispersion model were combined with daily emission factors derived from logistic growth models to construct a taxon-specific predictive index. This index was then incorporated into machine learning models alongside additional geospatial predictors to jointly estimate daily pollen counts by taxon. Model performance was evaluated using daily pollen counts measured by automated sensors at multiple sampling sites across Atlanta, Georgia.

**Figure 1.**
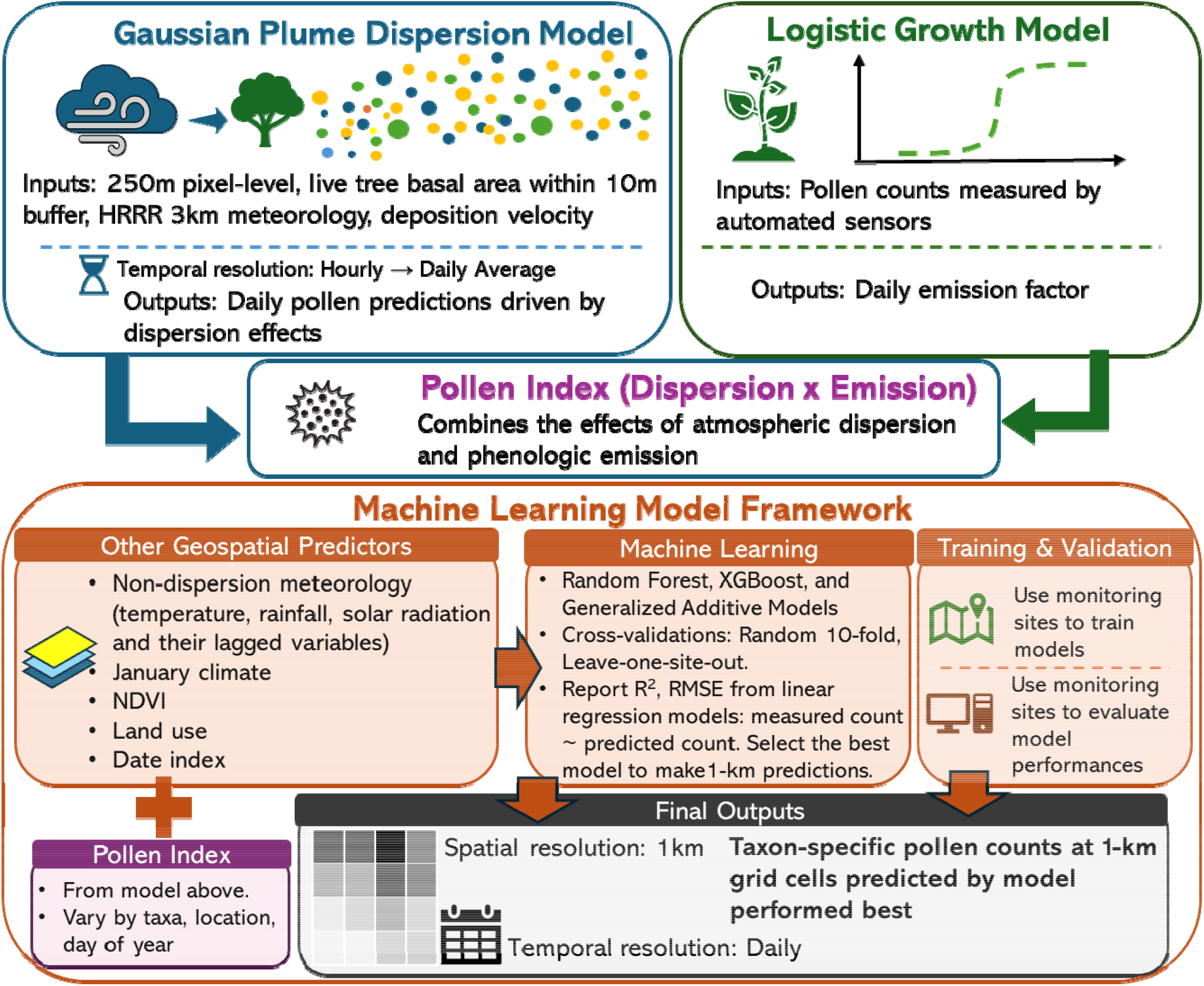
Overview of the hybrid modeling framework for estimating daily taxon-specific pollen concentrations at 1-km resolution.

### Study Area and Pollen Measurement Data

The study area included five counties in the Atlanta metropolitan region (Cobb, Clayton, Gwinnett, Fulton, and DeKalb), which encompasses an area of 4,484 square kilometers. Figure 2 shows the locations of pollen sampling sites, a PollenSense (PollenSense LLC) automated pollen sensor, and sample raw images returned by the device. The PollenSense sensors consist of an air pump, a sampling tape, a high-resolution camera that captures images of pollen particles collected on the tape, and a built-in Wi-Fi router that transmits these images to the company’s cloud platform for automated classification. Pollen grains are identified to the taxonomic level and enumerated using an AI-based image recognition algorithm. Images are typically captured at a rate of 1 to 3 per minute, and the smallest time unit of raw pollen count data provided by PollenSense is at an hourly temporal resolution.

**Figure 2.**
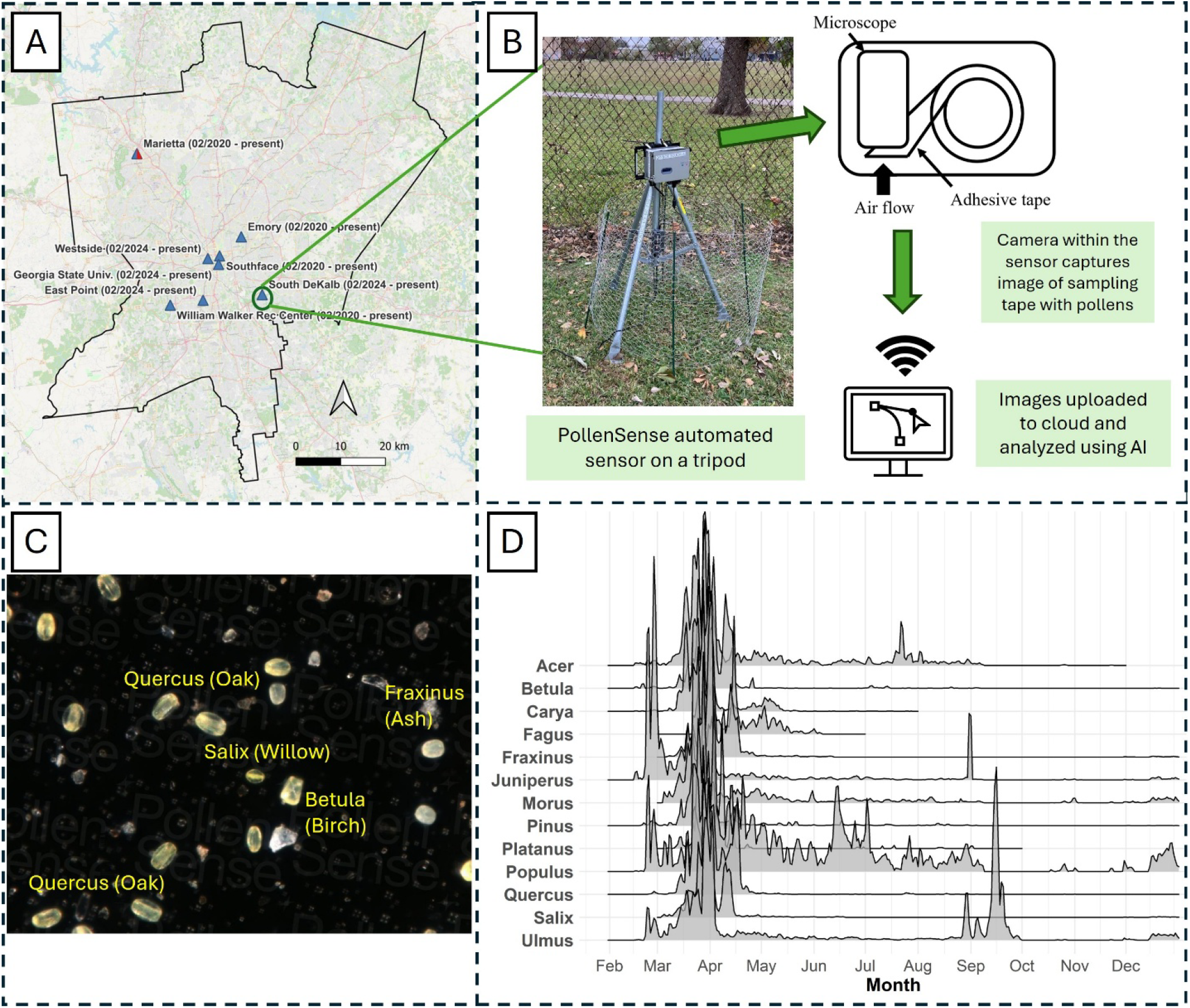
(A) Study region and locations of pollen monitoring sites, (B) schematic of the PollenSense automated sensor, (C) an example image took by sensor, and (D) temporal variation in pollen measurements collected by the sensors in 2024.

The deployment^33^ of PollenSense sensors in Atlanta began in 2020 with data collected in 2020 and 2021 comprising pilot monitoring efforts at three sampling sites (Emory, Southface, and Marietta). A full and standardized sampling protocol was implemented in 2024. Additionally, the number of sites and sensors increased to eight sites and nine total sensors in 2024. Most sites (East Point, Georgia State University, Marietta, South DeKalb, Southface and William Walker Recreation Center) were equipped with a single sensor, while one site (Emory) was equipped with two sensors. Notably, the Marietta site was co-located with Atlanta’s NAB station. A comparison of pollen data obtained from the automated sensors and the NAB method has been previously published.^33^

### Predictive Model Development

Taxon-specific pollen predictive models were developed based on machine learning fusing data curated from the Gaussian plume dispersion model, phenology model, and other geospatial data considered predictors of pollen. We followed a general framework commonly used in air pollution exposure modeling,^34, 35^ in which the model-fitting dataset consists of observed pollen measurements paired with predictors generated at the locations of monitoring sites. After model training, the fitted relationships were applied to generate predictions at locations without pollen measurements using corresponding predictor data. The preparation of machine learning predictors is described below.

#### Dispersion modeling

We chose Gaussian plume dispersion model to characterize the spatial and temporal variation in pollen transport under local meteorology conditions. The strength of Gaussian plume dispersion model included providing high-resolution spatiotemporal concentration gradients and a computationally efficient, scalable framework for modeling large geographic domains. Although Gaussian plume models have been primarily designed for modeling the near-surface transport of airborne particles, it has also been used in modeling bioaerosols.^36, 37^

In this study, we revised the algorithm of a neighborhood-scale Gaussian plume dispersion model to^38^ adapt for pollen. First, particle’s dispersion trajectory varied by particle size, density, and surface-to-volume ratio, we modified the standard Gaussian plume model by incorporating a deposition function that allows specification of taxon-specific deposition velocities (m s□^1^). The revised algorithm, along with implementation examples, is provided in Supplementary Document Section S1.

Second, the source inputs of our revised Gaussian plume dispersion model were prepared using the Live Tree Species Basal Area dataset^39^ published by the U.S. Department of Agriculture Forest Service (USFS). This dataset contains the basal area of specific tree species at a 250-m spatial resolution for 324 species across the U.S. We considered all 250-m grid cells within a 10-km buffer area as the site’s pollen sources.

For each 250-m grid cell, we selected and grouped species into genera that detected in the pollen measurements from automated sensors. During our sampling periods from 2020-2021 and 2024, pollen from 17 distinct tree genera were detected (Supplementary Table S2): *Acer, Alnus, Betula, Carya, Fagus, Fraxinus, Celtis, Juniperus, Morus, Olea, Pinus, Platanus, Populus, Quercus, Salix, Ulmus*, and *Juglans*. Among these genera, *Alnus* and *Olea* were not present in the Live Tree Species Basal Area dataset within a 10-km buffer of the sampling sites; *Celtis* and *Juglans* were detected at only a single sampling site with very low pollen counts; and *Fagus, Platanus*, and *Juglans* were not supported in earlier versions of the PollenSense AI classification algorithm, thereby they were unavailable in the 2020–2021 sampling data. As a result, our primary analyses focused on 11 genera (*Acer, Betula, Carya, Fraxinus, Juniperus, Morus, Pinus, Populus, Quercus, Salix*, and *Ulmus*) across the five-year study period, with two additional taxa (*Fagus* and *Platanus*) included in the 2024 study year only.

Notably, the USFS’s Live Tree Species Basal Area dataset was developed based on observations between 2000 and 2009 as recorded by the Forest Inventory and Analysis (FIA) program, preceding our study period by approximately 10 to 20 years. To adjust for the change of vegetation coverage, we used recent U.S. Geological Survey (USGS) tree canopy data^40^ to scale the basal area values. We obtained annual tree canopy cover raster files, which represent the percentage of land area covered by tree canopy within each 30-m grid cell across the continental U.S., for 2000 and 2020 through 2023. As 2023 was the latest year available at the time of analysis, 2023 data was used for both 2023 and 2024 study years.

We extracted all USGS 30-m grid cells and aggregated the percent canopy cover within each 250-m grid cell corresponding to the Live Tree Species Basal Area dataset. We then calculated a 250-m cell-specific ratio of tree canopy cover between study year and 2000, the year that USFS dataset was first compiled. For example, the ratios of “% tree canopy cover in 2023/ % tree canopy cover in 2000” were used to multiply the USFS basal area values to generate updated basal values for study year 2023-2024. The adjusted 250-m resolution basal area data were then used to represent taxon-specific pollen emission sources from each 250m grid cell.

Third, meteorological inputs to the Gaussian plume dispersion model were derived from the High-Resolution Rapid Refresh (HRRR) model based on our recently developed method.^41^ HRRR provides meteorological fields at 3-km spatial resolution and hourly temporal resolution. Correspondingly, our Gaussian plume model produced hourly estimates of pollen levels at receptor locations, driven by the pollen emission sources within 10km buffers and the meteorology conditions provided by the nearest 3-km HRRR. During model training, receptor locations were pollen sampling sites, whereas during prediction the receptor locations were defined as 1-km grid cells across the Atlanta metropolitan area. Because Gaussian plume dispersion models can generate unrealistically large estimates under highly stable atmospheric conditions, hourly estimates were aggregated to daily averages and then natural log transformed to reduce the influence of extreme values.

#### Phenological modeling

We designed a two-layer logistic growth model^42^ to estimate seasonal variation in daily pollen counts. Logistic growth models have been widely used in ecology, agriculture, and infectious disease modeling, as biological growth curves.^42^ Thus, logistic growth models are commonly used in predicting plant growth,^43, 44^ including in several pollen studies^45, 46^ to predict the seasonal variation of pollen. We fit a logistic growth model with equation (1) for each pollen taxon and each study year separately:

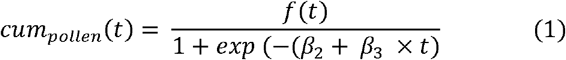

where *cum*_*pollen*_ (*t*) is the cumulative pollen count by the *t*^*th*^ day of the pollen season, *β*_2_ is an intercept-like term, *β*_3_ is the slope representing the growth rate, and *f*(*t*) is the time-varying carrying capacity as trees gradually “wake up” before producing pollen. *f*(*t*) is modeled as another logistic function as equation (2):

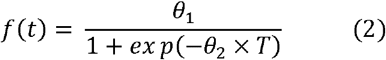

where *θ*_1_ is the overall carrying capacity calculated as the maximum value of cumulative pollen during a pollen season and *T* is the growing period of tree taxa, assumed to end at the middle of the pollen season.

We simulated the *β*_2_, *β*_3_, *θ*_1_, *θ*_2_ using nonlinear least squares regression models built with equations described above. In this study, each year’s pollen season was defined as the period between the first and last dates on which pollen was detected. For some taxa, low levels of pollen were occasionally detected during fall or winter. Because these low-level observations did not support stable growth curve estimation using a logistic model, the end of the pollen season was restricted to no later than the 150^th^ day of the year for all taxa except *Ulmus. Ulmus* are widely known to have a secondary pollen season in the fall and therefore a separate logistic growth model was fitted for the fall season.

#### Taxon-specific predictive indexes

In the next step, the estimated logistic growth parameters were converted into a daily growth rate metric by calculating the proportion of each day’s growth relative to the maximum carrying capacity. This metric will serve as an adjustment factor to model daily variability in pollen emissions when applied to the daily average of Gaussian plume dispersion model outputs. The productivity of taxa-specific “daily emission factor x natural log transformed daily average Gaussian plume model outputs” estimates represented pollen predictive indexes that were used as a predictor in taxa-specific machine learning models.

#### Other predictors

In addition to the taxon-specific indices derived from the dispersion and logistic growth models, the relationship between these indices and observed pollen counts may be influenced by local landscape characteristics and meteorological factors not directly related to dispersion, such as precipitation. To account for these effects and better capture the spatial and temporal variations in pollen counts, we applied machine learning models incorporating a comprehensive set of meteorological and land-use predictors. The full list of predictors is provided in Supplementary Table S2.

### Machine Learning Model Development and Evaluation

We fitted Random Forest (RF) and Extreme Gradient Boosting (XGB) models with predictors described above, validated by daily pollen counts measured by automated sensors. During model fitting, RF and XGB models were trained separately for each pollen taxon, and their predictions were combined in a Generalized Additive Model (GAM) for ensemble estimates. The GAM incorporated smooth functions of RF predictions, XGB predictions, and Julian day to capture both nonlinear relationships and seasonal trends. The model (RF, XGB, or GAM) that demonstrated the best predictive performance was subsequently applied to generate daily taxon-specific pollen predictions on 1-km grid cells by leveraging the learned relationships and gridded predictors at each 1-km cell.

During model fitting, model performance was evaluated using both random 10-fold cross-validation (CV) procedures and leave-one-site-out CV (LOSO-CV). In the random 10-fold CV, the full study dataset was randomly divided into 80% for model training, 10% for testing of RF and XGB models, and the remaining 10% for testing the GAM ensemble. Each model was trained on 80% of the data and used to predict pollen counts for a separate 10% holdout testing set. The GAM ensemble was trained using out-of-sample predictions from the RF and XGB models and validated against the independent ensemble testing set. This process was repeated ten times such that each site was included in the testing set once. For each iteration, we calculated the coefficient of determination (R^2^) and root mean squared error (RMSE) between measured and predicted pollen counts and reported the average R^2^ and RMSE across all folds. RMSE values were normalized by taxon-specific peak counts to show a relative magnitude of error.

We further evaluated model performance in capturing both temporal and spatial variability in pollen counts. Temporal prediction performance metrics (R^2^ and RMSE) were computed based on deviations between daily predictions and annual mean concentrations at each site. Spatial prediction performance was typically assessed using long-term average concentrations by site. Because the number of pollen sampling sites (n = 8) was small and insufficient to support a site-average regression, we employed a LOSO-CV approach to evaluate spatial prediction accuracy. During LOSO-CV, models were trained using data from seven sites and validated on the remaining site. This process was performed eight times so that each site had served as the validation site once.

### Sensitivity Analysis

To assess potential overfitting and evaluate the robustness of our modeling framework, we implemented two sensitivity analyses.

First, we conducted a cross-year calculation of the pollen growth rate. Logistic growth parameters were estimated using the growth rates from 2024’s models and 2021’s pollen season starting dates. These parameters were then applied to predicted pollen counts for 2021. Second, we developed separate models for the pollen season and non-season periods. We assume that pollen emission rates vary minimally outside the primary pollen season and that observed concentrations during the non-season are primarily driven by atmospheric dispersion processes. Therefore, the pollen index for the non-season model was estimated solely using Gaussian plume dispersion model derived from HRRR meteorological inputs and took a natural log transformation.

This study used QGIS 3.36.1 for processing geospatial predictors and R 4.3.3 for model development. Multiple R packages were utilized throughout the analysis. The parameters of the two-layer logistic growth model were estimated using the nlsLM() function from the minpack.lm package. Machine learning models were implemented using the randomForest, xgboost, and mgcv packages for RF, XGB, and GAM, respectively. Both R and QGIS were used in visualization of predicted taxon-specific pollen at 1-km grid cells across study region.

## Results

### Automated Sensor Pollen Measurements

**Figure 2** presents the locations of eight pollen monitoring sites in the five-county area of Atlanta, GA (Panel A), the configuration of the PollenSense automated sensors (Panel B), an example image took by sensor with annotations of pollen taxa (Panel C), and temporal variation in pollen measurements collected by the sensors in 2024 (Panel D). The temporal variation of study tree taxa for other two study years (2020 and 2021) is available in supplementary document Figure S1. To account for large differences in pollen abundance across taxa, daily counts were standardized by their standard deviations. The peak values of pollen count and pollen season for each of the 13 study taxa are available in supplementary document **Tables S3-5**.

Taxon-specific daily pollen emission factors estimated from logistics growth models are shown in Supplementary **Figure S2**. Without completed full season sampling, emission factors for 2022 and 2023 were derived using the observed pollen season start dates for each respective year and the growth rate parameters (carrying capacity, growth rate of pollen release, growth rate of trees “wake up”) estimated from 2024. As a result, the daily emission factor curves for these three years exhibit similar temporal shapes but are shifted in time according to differences in season onset.

### Model Performance

**Figure 3** presents the results of RF, XGB, and ensemble GAM models. The three models demonstrated comparable predictive performance, with XGB generally achieving slightly higher R^2^ values than the other approaches (**Figure 3** Panel A). Model performance varied substantially across pollen taxa and study year and no consistent pattern emerged indicating that any specific taxa or single year consistently achieved higher R^2^ values. Overall, *Betula* and *Quercus* exhibited higher predictive performance than other taxa, with R^2^ values ranging from 0.69 to 0.92 for *Betula* and from 0.71 to 0.89 for *Quercus*. The R^2^ visualized in Panel A of **Figure 2** are provided in Supplementary **Tables S6–S8**. The R^2^ for non-season were also calculated and provided in Supplementary **Tables S9-11**. Because non-season pollen counts were predominantly zero, model performance during the non-season was generally low, as reflected by lower R^2^ values comparing to the pollen season.

**Figure 3.**
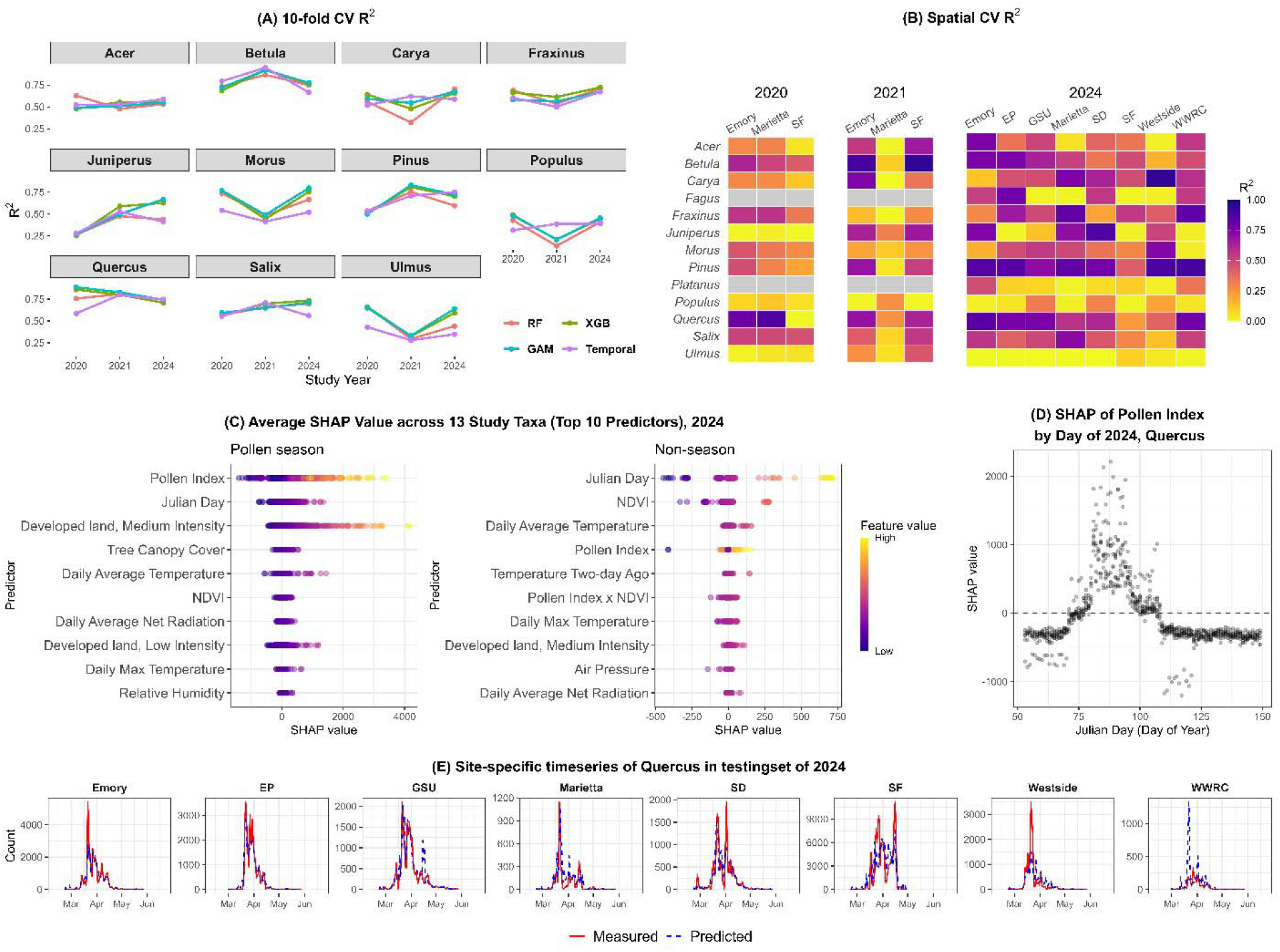
(A) Ten-fold cross-validation (CV) R^2^ by taxa, model year, and modeling approaches, Random Forest (RF), Extreme Gradient Boosting (XGB), Generalized Additive Models (GAM), and a temporal R^2^ from XGB across 11 tree taxa for three sampling years (2020, 2021, and 2024). *Fagus* and *Platanus* were not included here because these two taxa were only analyzed for 2024. (B) Site-specific R^2^ from leave-one-site-out (LOSO) CV for each tree taxon across monitoring sites in 2020, 2021, and 2024. (C) SHAP values for the top 10 predictors identified through the importance averaged across 13 XGB-modeled tree taxa in 2024, separate by pollen season and non-season. In this study, pollen season was defined as between the first date and last date with pollen detected, while the last date is no later than the 150^th^ day of year. *Ulmus* have two seasons in a year. (D) SHAP dependence plot for the pollen index versus Julian day for *Quercus* in 2024. (E) Timeseries plots of XGB-predicted *Quercus* pollen count during pollen season among the testing sets from 10-fold random cross validation approach for 2024. Figures for 13 study taxa and sampling years are available in Supplementary Figures S3-S4. Abbreviations: Emory (Emory University), EP (East Point), GSU (Georgia State University), SD (South DeKalb), SF (Southface), and WWRC (William Walker Recreation Center).

**Figure 4.**
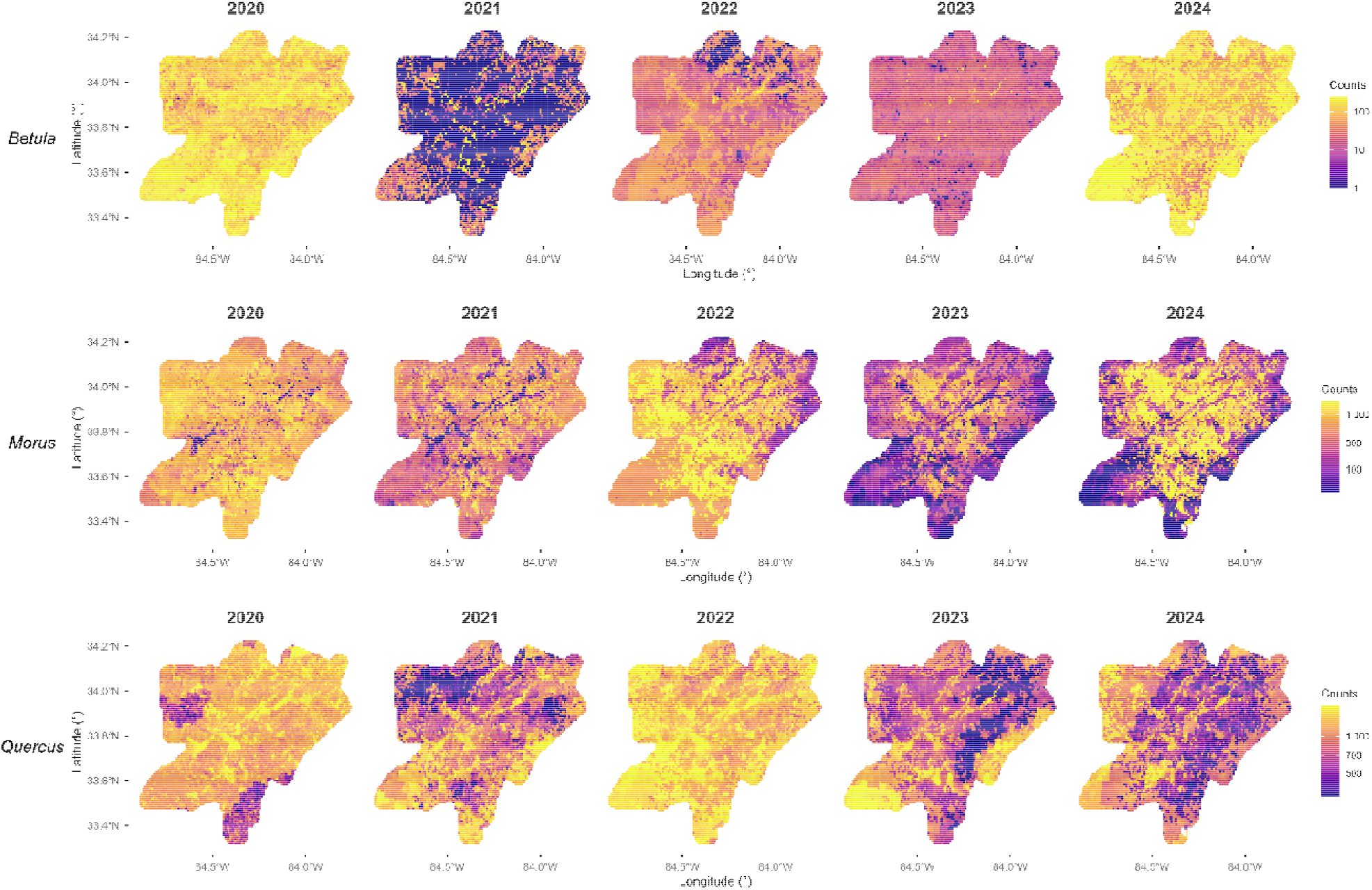
Predicted pollen at 1-km resolution for three allergenic taxa, *Betula, Morus*, and *Quercus*, on March 30 from 2020 to 2024, across the five-county area of Atlanta, GA. The warm color shows higher counts and cold color shows lower counts.

Given that XGB models generally performed better than RF and GAM, we then used the XGB to generate the 1-km predictions. Prior to that, we evaluated the influence of individual predictors on pollen predictions using calculated SHapley Additive exPlanations (SHAP) values for individual predictors from XGB models. **Figure 3** Panel C presents SHAP values for the top 10 predictors ranked by their relative importance. For the pollen season, the pollen index was the most important predictor across taxa. **Figure 3** Panel D illustrates this relationship using the 2024 *Quercus* model as an example. That is, the pollen index was negatively associated with pollen counts before and after the seasonal peak but positively associated during the peak period. This pattern reflects the combined effects of atmospheric dispersion and phenology. **Figure 3** Panel E shows the time-series of *Quercus* for the 2024 pollen season, including the time-series plots for both measured and XGB-predicted counts. The timeseries for all 13 study taxa and study years are available in supplementary document Figure S3-S4.

### Sensitivity Analysis

The 10-fold CV R^2^ values for the cross-year models and their comparisons with the 2021 R^2^ values from the primary analysis are presented in Supplementary **Table S12**. Cross-year models exhibited similar R^2^ values and RMSEs as models trained using same-year observations. The R^2^ values achieved by the cross-year models indicate that the modeling framework remains robust for most taxa. The cross-year R^2^s of *Betula* (0.76-0.90) and *Quercus* (0.86-0.93) were comparable to the main analysis. Accordingly, we applied the cross-year emission factor strategy to generate daily pollen predictions for 2022 and 2023, two years with insufficient measurement data to fit logistic growth models.

## Discussion

Leveraging a multi-site sampling network developed based on novel automated pollen sensors and a high-resolution geospatial modeling framework, we developed a method to predict taxon-specific pollen counts at a 1-km spatial and daily temporal resolution. Model performance was comparable to that reported in previous studies applying machine learning approaches.^29^ Our framework provides several important advances in pollen predictions. First, it captures both temporal and spatial variation in pollen. The 1-km spatial resolution provides fine-scale estimates that characterize neighborhood-level variability in pollen exposure, which is particularly relevant for assessing individual-level exposure; the continuous daily predictions across through year can help epidemiologic studies to examine the lagged effects of pollen. Second, the framework is scalable and leverages widely available, nationwide datasets, allowing extension across years and geographic regions.

In comparison of studies^18, 24, 28, 29^ using NAB data to develop predictive models, the multi-year, multi-site pollen measurement data from automated sensors provide more representative reference data for the spatial and temporal variation of pollen counts within a city. According to the pollen counts measured by our monitoring network, there is considerable variability in pollen abundance across study years and monitoring sites (**Tables S3-4**). Every year, *Quercus* had the highest pollen counts, followed by *Pinus*. This finding is consistent with previous studies (Lo et al.^29^ and Lappe et al.^6^) that analyzed Atlanta’s NAB pollen data. In terms of time of observed pollen season, in 2024, the first detections of pollen generally occurred in mid-February, which is comparable to pollen season reported in Lo et al.^29^ but earlier than reported in Lappe et al.^6^ This discrepancy likely reflects the change in pollen season over years. While both prior studies reported long-term averages, Lo et al.^29^ focused on more recent years (2003–2017) where Lappe et al.^6^ included more historical period (1993–2018). Earlier pollen season onset was observed in more recent years and is correlated with increasing temperatures.^47^

Our models demonstrated strong predictive performance, with particularly high R^2^ values for *Betula* and *Quercus*, two taxa commonly associated with allergic responses. Analyses conducted via pollen models developed by other groups, such as the PlantScope time series model,^30^ PECM,^24^ and CMAQ/HYSPLIT system,^25-27^ were not specifically focused on Atlanta or have different temporal resolution. Thus, direct comparisons are not appropriate. Instead, we compared our R^2^ to the data-driven machine learning models developed by Lo et al.^29^ for Atlanta, GA (R^2^=0.77 for *Quercus*) and found comparable results.

We observed a strong influence of pollen abundance on model performance as taxa with lower pollen counts generally yielded lower R^2^ values. This pattern is consistent with findings from air pollution modeling studies, in which reduced ambient concentrations are often associated with reduced predictive performance. An advantage of using multi-year data in this study is that, for individual taxa, we are able to compare each year’s R^2^ to the year’s pollen abundance. For example, in 2020 *Juniperus* showed substantially weaker predictive performance (R^2^ = 0.27 from XGB) compared with other years (R^2^ = 0.59 for 2021, R^2^ = 0.63 for 2024). That same year, *Juniperus* pollen measurements were also substantially lower (site-specific peak count = 12.55) than in subsequent years (site-specific peak counts = 5057.05 in 2021 and 1354.84 in 2024). Additionally, this pattern explains the low R^2^ values of non-season models where pollen counts are generally low.

Our sensitivity analysis comparing same-year and cross-year models indicated generally similar modeling performance. This finding suggested a robust estimate of 2024’s emission factor, which may be benefited from the relatively larger number of monitoring sites in this year. The expanded network provided broader spatial coverage across Atlanta and yielded more representative emission factors. The robust modeling performance also suggested the potential for developing a forecasting application of our model, that the seasonality patterns derived from one year could be used for predicting the seasonality in subsequent years. By examining the change of performance, performance reductions were relatively greater for *Pinus*, with R^2^ reduced by more than 30%. The different seasonal patterns across years (Supplementary **Figure S1**) help explain these discrepancies. For example, *Pinus* exhibited a substantially higher peak in 2024 compared with 2021, whereas the emission factors were derived from single-year data and therefore do not capture interannual variation in carrying capacity. Works on developing forecast system should pay attention to the estimation of carrying capacity to improve the cross-year predictions.

In this study, we advanced the standard Gaussian plume dispersion model by incorporating several innovations, including the add-on function of taxon-specific deposition velocities, taxon-resolved source inputs, and spatially resolved meteorological data. Adding deposition velocity into the Gaussian plume model increases the relative influence of nearby sources and also increased the R^2^ for most study taxa except for *Betula* and *Salix. Betula* pollen is known to undergo long-distance atmospheric transport, a phenomenon that has been well documented in previous studies.^48, 49^ The long-range transport of *Salix* pollen has been less studied, but the catkin structure of *Salix* flowers may facilitate wind-floated dispersion over longer distances. Also due to this, we selected relatively smaller deposition velocities for these two taxa when multiple values were available in literature. This revised Gaussian plume model with a function of setting taxon□specific deposition velocities is applicable to air pollution studies, given that different air pollutants also have different aerodynamic features.

This study has several important strengths. First, the two-layer logistic growth model we developed seems not affected by the incomplete sampling schedule. For some taxa, such as *Fagus* in 2024, *Juniperus*, and *Betula* in 2021, the estimated emission curves (Figure S2) appeared truncated. This was due to the start of sampling is later than the start of pollen season of those taxa, which did not fully capture the full pollen season (Table S5). Nevertheless, the intercept term in the logistic growth model accounts for this truncation.

Second, the machine learning models helped address gaps between pollen dispersion processes and measurement data by incorporating additional meteorology or land use variables. For example, rainfall can have both immediate and delayed impacts on pollen: recent precipitation can reduce airborne pollen through washout, whereas rainfall in several days ago may stimulate trees to produce more pollen. To capture these relationships, we incorporated meteorological predictors with one and two days prior, which have been shown to be important predictors of daily pollen levels.

Our modeling framework possesses some limitations, especially in the method of calculating the taxon-specific index. On one hand, in this study, we did not fit site-specific logistic growth models because 1-km predictions cannot be generated based on its location related to sampling sites that provide daily growth curves. Thereby, the daily pollen emission factors were estimated using data combined across all study sites to represent the aggregate of the five-county Atlanta area. However, pollen phenology can vary substantially at fine spatial scales due to microclimate variations in temperature and solar radiation. Future work will focus on developing more sophisticated phenology models that incorporate spatiotemporal variability in pollen release.

On the other hand, the USFS basal area data used in this study was developed more than a decade ago. Although we adjusted these data using more recent tree canopy information, such adjustments were not taxon-specific. In addition, the 250-m spatial resolution of the basal area data does not capture the locations of individual trees. Future improvements could leverage high-resolution satellite imagery to generate up-to-date, tree-level information, which would enhance the accuracy of source characterization.

Overall, this is the first study to provide daily pollen counts at a neighborhood-level (1km grid cell). The generated data can help identify the predominant pollen taxa responsible for pollen allergy. Moreover, the continuous daily time series of taxon-specific predictions spanning five years provides gapless exposure metrics that can be readily incorporated into epidemiologic analyses.

## Data and Code Availability

The revised Gaussian plume dispersion model source code and instructions is publicly available on GitHub (see Supplementary Document Section S1). Predicted pollen counts at 1-km spatial resolution during the pollen season of 2024 are visualized an interactive R Shiny application (https://xueying-zhang.shinyapps.io/AtlantaPollenMap). The full dataset of predicted pollen counts is available from the corresponding author upon request.

## Acknowledgements

This work was supported by the National Institutes of Health (NIH) under grants P30ES030285 and P50MD015496, and by the National Aeronautics and Space Administration (NASA) under grant 80NSSC26K0265 (24-HAQ24-0043).

