## Supporting Information for "Estimating Daily Taxon-specific Tree Pollen at a 1-km Resolution in Atlanta, GA from 2020 to 2024"

Corresponding author:

**Section S1. Revised Gaussian plume dispersion model with deposition**

The Gaussian plume dispersion model used in this study was adapted from the Research LINE-source dispersion model (R-LINE; Snyder et al. 2013), which was originally developed for near-surface, traffic-related air pollution. Because pollen grains differ substantially from typical air pollutants in terms of particle size, density, and surface-to-volume ratio, we modified the standard formulation to incorporate taxon-specific dry deposition processes.

The modification is centered on deposition velocities (the unit is m s⁻¹) represent the gravitational settling of pollen particles during atmospheric transport. Higher deposition velocities indicate faster settling and shorter transport distances. In the Gaussian plume framework, this results in relatively greater contributions from nearby pollen sources to the estimated pollen concentrations at receptor locations.

The revised Gaussian plume model is expressed as:

$$C=\frac{Q}{\sqrt{2\pi} U \sigma_{z}} exp\left( - \frac{(z -h)^{2}}{2\sigma_{z}^{2}} \right) exp\left( - \frac{v_{d} x}{UH_{mix}} \right)$$

Where

- $C$ was designed to be in a mass/volume unit (e.g., ug/m^3^) if the input emission factor ($Q$) is in the unit of gram/meter/second. Since our input (in unit of basal area in sqm^2^) is not the required unit of R-LINE, then in our study, $C$ is unit less.
- $Q$ is the daily emission factor of pollen, proxy as tree basal area
- $U$ is wind speed (m/s)
- $\sigma_{z}$ is a parameter (m/s) indicating atmospheric stability estimated based on surface friction velocity (m/s) and convective velocity (m/s).
- $z$ is receptor height and we set it to ground level (zero meter).
- $h$ is effective release height and we set it to 10 meters.
- $v_{d}$ is the dry deposition velocity of pollen taxa (m/s)
- $x$ is the downwind distance between source and receptor.
- $H_{mix}$is atmospheric mixing height (m).

Model implementation and user interface: In addition to revising the dispersion algorithm, we modified the model interface to allow users to specify taxon-specific deposition velocities. The revised user interface includes an option to enable or disable dry deposition:

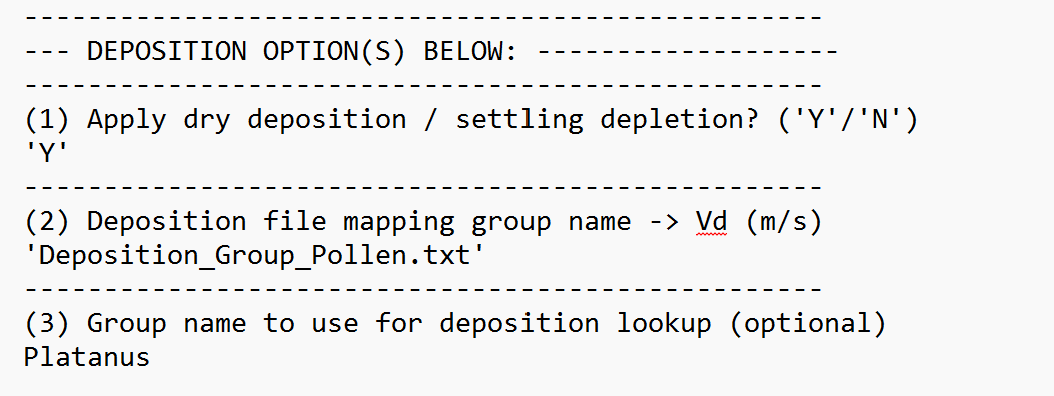

If answered “Y” for the question (1) Apply dry deposition / settling depletion (‘Y’/’N’), then the deposition function is enable, users are prompted to another file (“Deposition_Group_Pollen.txt” in this example) for storing deposition velocities. This file provided taxon-specific deposition parameters:

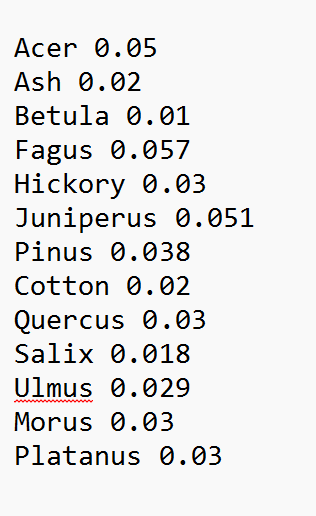

For this study, we use deposition velocities found in literature. The number and corresponding literature references are available in the following Table S1. For four genera (*Acer, Carya, Morus, Platanus*) for which deposition velocities were not available, we assigned an average value of 0.03 m s⁻¹ based on the mean of values reported for other taxa.

| Table S1. Deposition velocities (m/s) used in Gaussian plume dispersion model. The number in brackets is the deposition velocity reported in each literature. | | | |
| --- | --- | --- | --- |
| **Taxon** | **Common name** | **Deposition velocity (m/s)** | **Source** |
| *Acer* | Maple | 0.03 | Not found.^a^ |
| *Alnus* | Alder | 0.021 | Eisenhut 1961 |
| *Betula* | Birch | 0.01 | Eisenhut 1961 (0.022); Sofiev et al. 2006 (0.012); Li et al. 2015 (0.019) ; Zhang et al. 2015 (0.031) |
| *Carya* | Hickory | 0.03 | Not found.^a^ |
| *Fagus* | Beech | 0.057 | Eisenhut 1961 |
| *Fraxinus* | Ash | 0.02 | Eisenhut 1961 (0.022); Li et al. 2015 (0.017) ; Zhang et al. 2015 (0.021) |
| *Juniperus* | Juniper | 0.051 | Borrell 2012 (0.07); Fang et al. 2018 (0.035) |
| *Morus* | Mulberry | 0.03 | Not found.^a^ |
| *Olea* | Oliver | 0.032 | Zhang et al. 2015 (0.032) |
| *Pinus* | Pine | 0.038 | Lorenz and Murphy 1989 (0.031); Eisenhut 1961 (0.031); Li et al. 2015 (0.039) ; Dyakowska et al. 1937 (0.04) Fang (0.046) ; Zhang et al. 2015 (0.042) |
| *Platanus* | Plane | 0.03 | No direct number, but the relative deposition rate for Platanus were mentioned in Adams-Groom et al. 2017. |
| *Populus* | Poplar, Cottonwood | 0.02 | Xu et al. 2012 |
| *Quercus* | Oak | 0.03 | Eisenhut 1961 (0.035) ; Li et al. 2015 (0.018) ; Fang et al (0.016) ; Zhang et al. 2015 (0.039) ; Xu et al. 2012 (0.042) |
| *Salix* | Willow | 0.018 | Zhang et al. 2015 (0.018) ; Gregory et al. 1973 (0.022) |
| *Ulmus* | Elm | 0.029 | Gregory et al. 1961 (0.032); Li et al. 2015 (0.022) ; Zhang et al. 2015 (0.034) |
| *Juglans* | Walnut | 0.034 | Bodmer, 1922 (0.037); Li et al. 2015 (0.030) |
| Note: a. For taxa whose deposition velocities did not find in literature, assign value based on the pollen shape, size, and clearance comparing to other taxa with reported deposition velocities. | | | |

R-LINE’s temporal resolution is determined by the temporal resolution of meteorological inputs. In practice, users typically run the model at an hourly resolution because shorter time units (e.g., 1 minute) may not enough for the full plume to develop, while longer time units (e.g., daily averages) do not produce physically meaningful “average” concentration predictions based on the daily average wind speed or wind direction. In this study, we run R-LINE using the hourly, 3-km resolved High-Resolution Rapid Refresh (HRRR) meteorology data.

The full implementation is written in Fortran, as same as used by the standard R-LINE model. The source code of the revised model is publicly available on GitHub (https://github.com/zxy1219/RLINE-deposition).

**Section 2. Supplementary document data for study data and modeling results.**

| Table S2. Predictor variables used in the machine learning models | | | | | |
| --- | --- | --- | --- | --- | --- |
| **Category** | **Predictor** | **Description** | **Source** | **Temporal Resolution** | **Lag Structure / Notes** |
| Process-based | Taxon-specific pollen index | Dispersion × emission derived from Gaussian plume + logistic growth models | This study derived | Daily | Same day |
| Meteorology (HRRR) | Temperature | Mean, maximum, and diurnal range | HRRR | Hourly → Daily | Lag 0–2 days |
|  | Relative humidity | Mean relative humidity | HRRR | Hourly → Daily | Lag 0–2 days |
|  | Surface pressure | Mean and daily range | HRRR | Hourly → Daily | Same day |
|  | Wind | Maximum surface wind gust | HRRR | Hourly → Daily | Same day |
|  | Mean sea-level pressure | Atmospheric pressure at sea level | HRRR | Hourly → Daily | Same day |
|  | Ground heat flux | Surface energy exchange | HRRR | Hourly → Daily | Same day |
|  | Net radiation | Surface net radiation | HRRR | Hourly → Daily | Same day |
|  | Cloud cover | Mean cloud fraction | HRRR | Hourly → Daily | Same day |
|  | Solar radiation | Total beam solar radiation | HRRR | Hourly → Daily | Lag 0–2 days |
| Meteorology (Observed) | Precipitation | Daily precipitation | NOAA ISD (Atlanta airport) | Daily | Lag 0–2 days |
|  | Dew point | Daily dew point temperature | NOAA ISD | Daily | Lag 0–2 days |
| Seasonality | Julian day | Day of year to capture seasonal trends | Derived | Daily | Same day |
|  | January climate | January mean temperature, solar radiation, and total precipitation | HRRR / NOAA | Monthly | January of study year |
| Land use / Vegetation | NLCD classes | Area of each land-use type within 1-km grid | NLCD | Static | Spatial predictor |
|  | Tree canopy cover | Area of tree canopy within grid | NLCD / canopy dataset | Static | Spatial predictor |
|  | NDVI | Vegetation index representing greenness | VIIRS | 10-day | Assigned using nearest 10-day window |
| Notes: Observed precipitation from NOAA ISD was included to better capture extreme rainfall events not fully represented in HRRR outputs. | | | | | |

| Table S3. Peak daily pollen counts^a^ by taxon across sampling sites in 2024. | | | | | | | | | |
| --- | --- | --- | --- | --- | --- | --- | --- | --- | --- |
| **Taxon** | **Common name** | **Emory** | **EP** | **GSU** | **Marietta** | **SD** | **SF** | **Westside** | **WWRC** |
| *Acer* | Maple | 120.62 | 113.08 | 290.97 | 39.64 | 175.12 | 148.49 | 29.47 | 20.82 |
| *Alnus^b^* | Alder | 263.68 | 404.99 | 20.48 | 190.50 | 150.44 | 2764.89 | 574.11 | 24.65 |
| *Betula* | Birch | 481.57 | 470.80 | 92.24 | 42.58 | 234.17 | 4763.52 | 231.34 | 24.64 |
| *Carya* | Hickory | 173.42 | 260.79 | 265.46 | 877.50 | 774.36 | 4189.45 | 1432.26 | 14.84 |
| *Fagus* | Beech | 144.35 | 231.10 | 50.40 | 111.38 | 48.32 | 623.61 | 86.83 | 48.93 |
| *Fraxinus* | Ash | 264.40 | 199.08 | 134.37 | 276.07 | 299.99 | 2663.86 | 1259.04 | 26.60 |
| *Celtis* | Hackberry | 0 | 0 | 4.93 | 0 | 0 | 0 | 0 | 0 |
| *Juniperus* | Juniper | 339.27 | 39.29 | 327.44 | 218.19 | 1354.84 | 580.44 | 229.20 | 18.60 |
| *Morus* | Mulberry | 195.44 | 193.06 | 1342.71 | 1488.18 | 1402.58 | 487.00 | 5623.40 | 57.96 |
| *Olea^b^* | Oliver | 101.93 | 119.36 | 24.78 | 56.58 | 63.99 | 1015.84 | 75.33 | 12.80 |
| *Pinus* | Pine | 4061.66 | 3537.15 | 1765.17 | 1443.80 | 6002.63 | 9220.55 | 2199.40 | 325.99 |
| *Platanus* | Plane | 123.63 | 55.46 | 38.81 | 7.88 | 6.73 | 2857.56 | 11.90 | 9.88 |
| *Populus* | Poplar, Cottonwood | 149.65 | 91.15 | 56.76 | 54.23 | 66.62 | 38.25 | 70.69 | 13.80 |
| *Quercus* | Oak | 5398.84 | 3538.03 | 2100.60 | 1144.95 | 1949.61 | 11170.08 | 3493.17 | 331.67 |
| *Salix* | Willow | 300.22 | 260.86 | 117.89 | 1394.46 | 506.35 | 3795.69 | 5202.97 | 21.26 |
| *Ulmus* | Elm | 2587.46 | 326.42 | 1394.90 | 356.13 | 1226.68 | 43.94 | 204.39 | 5.93 |
| *Juglans^b^* | Walnut | 1.95 | 0 | 0 | 0 | 0 | 0 | 1.33 | 0 |
| ^a^Pollen counts returned by PollenSense automated sensors are not always integers because images captured during sampling intervals that span two consecutive hours are split between those hours. As a result, pollen grains identified in these images are proportionally allocated across hours, yielding non-integer counts.  ^b^*Alnus*, *Olea*, and *Juglans* are not included in the final model.  Abbreviations: Emory (Emory University), EP (East Point), GSU (Georgia State University), SD (South DeKalb), SF (South Face), and WWRC (William Walker Recreation Center) | | | | | | | | | |

| Table S4. Peak daily pollen counts^a^ by taxon^c^ across sampling sites in 2020 and 2021. | | | | | | | |
| --- | --- | --- | --- | --- | --- | --- | --- |
| **Taxon** | **Common name** | **2020** | | | **2021** | | |
|  |  | **Emory** | **Marietta** | **SF** | **Emory** | **Marietta** | **SF** |
| *Acer* | Maple | 47.97 | 5.43 | 232.40 | 47.96 | 228.83 | 360.4 |
| *Alnus^b^* | Alder | 127.03 | 10.87 | 250.54 | 237.09 | 38.43 | 1096.88 |
| *Betula* | Birch | 1431.81 | 1678.42 | 1635.97 | 78.36 | 1474.91 | 848.77 |
| *Carya* | Hickory | 595.86 | 3959.99 | 409.65 | 6.93 | 549.6 | 467.68 |
| *Fraxinus* | Ash | 180.60 | 42.20 | 108.74 | 36.69 | 577.12 | 138.41 |
| *Juniperus* | Juniper | 11.62 | 2.18 | 12.55 | 216.02 | 5057.05 | 2905.04 |
| *Morus* | Mulberry | 98.55 | 17.59 | 287.23 | 23.82 | 68.25 | 310.73 |
| *Olea^b^* | Oliver | 265.53 | 25.89 | 408.79 | 0 | 0 | 0 |
| *Pinus* | Pine | 1633.64 | 33.73 | 2334.81 | 373.68 | 6521.97 | 4057.19 |
| *Populus* | Poplar, Cottonwood | 41.54 | 9.29 | 88.52 | 17.76 | 3.44 | 160.78 |
| *Quercus* | Oak | 3812.39 | 28.11 | 2313.28 | 192.31 | 9486.3 | 8685.49 |
| *Salix* | Willow | 222.36 | 309.75 | 224.95 | 49.31 | 527.45 | 624.02 |
| *Ulmus* | Elm | 41.65 | 3.67 | 73.50 | 41.03 | 13.95 | 126.11 |
| ^a^Pollen counts returned by PollenSense automated sensors are not always integers because images captured during sampling intervals that span two consecutive hours are split between those hours. As a result, pollen grains identified in these images are proportionally allocated across hours, yielding non-integer counts.  ^b^*Alnus* and *Olea* are not included modeled.  ^c^No data is available for *Fagus*, *Platanus*, and *Juglans* in 2020 and 2021 as these taxa were not included in earlier versions of PollenSense’s AI.  Abbreviations: Emory (Emory University), EP (East Point), GSU (Georgia State University), SD (South DeKalb), SF (South Face), and WWRC (William Walker Recreation Center). | | | | | | | |

| Table S5. Taxon- and year-specific pollen season start and end dates, day of year (DOY), and season length. | | | | | | | | |
| --- | --- | --- | --- | --- | --- | --- | --- | --- |
|  |  |  | |  | | **Model Season Length (Days)** |  | |
|  |  | **Model Start** | | **Model End** | |  | **Last Non-zero Count** | |
| **Year** | **Tree** | **Date** | **DOY** | **Date** | **DOY** |  | **Date** | **DOY** |
| 2020 | *Acer* | 3/1/2020* | 61 | 5/10/2020 | 131 | 71 | 5/10/2020 | 131 |
| 2020 | *Alnus* | 3/1/2020* | 61 | 5/19/2020 | 140 | 80 | 5/19/2020 | 140 |
| 2020 | *Betula* | 3/1/2020* | 61 | 5/29/2020 | 150 | 90 | 6/30/2020 | 182 |
| 2020 | *Carya* | 3/5/2020 | 65 | 5/29/2020 | 150 | 87 | 6/30/2020 | 182 |
| 2020 | *Fraxinus* | 3/2/2020 | 62 | 5/29/2020 | 150 | 90 | 7/21/2020 | 203 |
| 2020 | *Juniperus* | 3/6/2020 | 66 | 5/4/2020 | 125 | 61 | 5/4/2020 | 125 |
| 2020 | *Morus* | 3/2/2020 | 62 | 5/29/2020 | 150 | 90 | 6/21/2020 | 173 |
| 2020 | *Pinus* | 3/2/2020 | 62 | 5/23/2020 | 144 | 84 | 5/23/2020 | 144 |
| 2020 | *Populus* | 3/6/2020 | 66 | 6/30/2020 | 182 | 118 | 10/29/2020 | 303 |
| 2020 | *Quercus* | 3/1/2020* | 61 | 5/29/2020 | 150 | 90 | 6/30/2020 | 182 |
| 2020 | *Salix* | 3/1/2020* | 61 | 5/29/2020 | 150 | 90 | 5/30/2020 | 151 |
| 2020 | *Ulmus* | 3/1/2020* | 61 | 5/27/2020 | 148 | 88 | 5/27/2020 | 148 |
| 2021 | *Acer* | 2/2/2021* | 33 | 4/30/2021 | 120 | 88 | 4/30/2021 | 120 |
| 2021 | *Alnus* | 2/2/2021* | 33 | 3/31/2021 | 90 | 58 | 3/31/2021 | 90 |
| 2021 | *Betula* | 2/3/2021 | 34 | 3/31/2021 | 90 | 57 | 3/31/2021 | 90 |
| 2021 | *Carya* | 4/1/2021 | 91 | 5/26/2021 | 146 | 56 | 6/14/2021 | 165 |
| 2021 | *Fraxinus* | 2/4/2021 | 35 | 5/27/2021 | 147 | 113 | 6/20/2021 | 171 |
| 2021 | *Juniperus* | 2/3/2021 | 34 | 4/30/2021 | 120 | 87 | 4/30/2021 | 120 |
| 2021 | *Morus* | 2/3/2021 | 34 | 5/27/2021 | 147 | 114 | 6/17/2021 | 168 |
| 2021 | *Pinus* | 2/3/2021 | 34 | 5/26/2021 | 146 | 113 | 6/20/2021 | 171 |
| 2021 | *Populus* | 2/10/2021 | 41 | 4/4/2021 | 94 | 54 | 4/4/2021 | 94 |
| 2021 | *Quercus* | 2/2/2021* | 33 | 5/26/2021 | 146 | 114 | 6/15/2021 | 166 |
| 2021 | *Salix* | 2/2/2021* | 33 | 4/30/2021 | 120 | 88 | 4/30/2021 | 120 |
| 2021 | *Ulmus* | 2/4/2021 | 35 | 4/28/2021 | 118 | 84 | 4/28/2021 | 118 |
| 2024 | *Acer* | 2/16/2024 | 47 | 5/29/2024 | 150 | 104 | 11/22/2024 | 327 |
| 2024 | *Alnus* | 2/12/2024 | 43 | 5/29/2024 | 150 | 108 | 12/30/2024 | 365 |
| 2024 | *Betula* | 2/16/2024 | 47 | 5/29/2024 | 150 | 104 | 12/29/2024 | 364 |
| 2024 | *Carya* | 2/16/2024 | 47 | 5/28/2024 | 149 | 103 | 7/28/2024 | 210 |
| 2024 | *Fagus* | 3/23/2024 | 83 | 5/29/2024 | 150 | 68 | 6/4/2024 | 156 |
| 2024 | *Fraxinus* | 3/5/2024 | 65 | 5/29/2024 | 150 | 86 | 12/31/2024 | 366 |
| 2024 | *Juniperus* | 2/16/2024 | 47 | 5/29/2024 | 150 | 104 | 12/31/2024 | 366 |
| 2024 | *Morus* | 3/3/2024 | 63 | 5/29/2024 | 150 | 88 | 12/31/2024 | 366 |
| 2024 | *Pinus* | 2/12/2024 | 43 | 5/29/2024 | 150 | 108 | 12/31/2024 | 366 |
| 2024 | *Platanus* | 3/6/2024 | 66 | 4/24/2024 | 115 | 50 | 9/1/2024 | 245 |
| 2024 | *Populus* | 2/16/2024 | 47 | 5/29/2024 | 150 | 104 | 12/31/2024 | 366 |
| 2024 | *Quercus* | 2/23/2024 | 54 | 5/29/2024 | 150 | 97 | 12/31/2024 | 366 |
| 2024 | *Salix* | 3/3/2024 | 63 | 5/29/2024 | 150 | 88 | 12/31/2024 | 366 |
| 2024 | *Ulmus* | 2/16/2024 | 47 | 5/29/2024 | 150 | 104 | 12/31/2024 | 366 |
| Note: For modeling purposes, pollen season was defined as the period between the earliest and latest dates on which nonzero pollen counts were detected. The end date was set to no later than day of year 150 so as to exclude sporadic detections outside the main pollen season. In 2020 and 2021, sampling began after the onset of some taxa’s pollen release; therefore, the model start date reflects either the beginning of sampling or the onset of the pollen season, whichever occurred later. Model start dates that reflect sampling start date are denoted with an asterisk (*). | | | | | | | | |

Figure S1. Temporal variation in tree pollen abundance measured by PollenSense sensors in Atlanta, GA across three years (2020, 2021, and 2024). Ridge plots show daily pollen counts aggregated across monitoring sites and scaled by one standard deviation within each year and tree taxon.

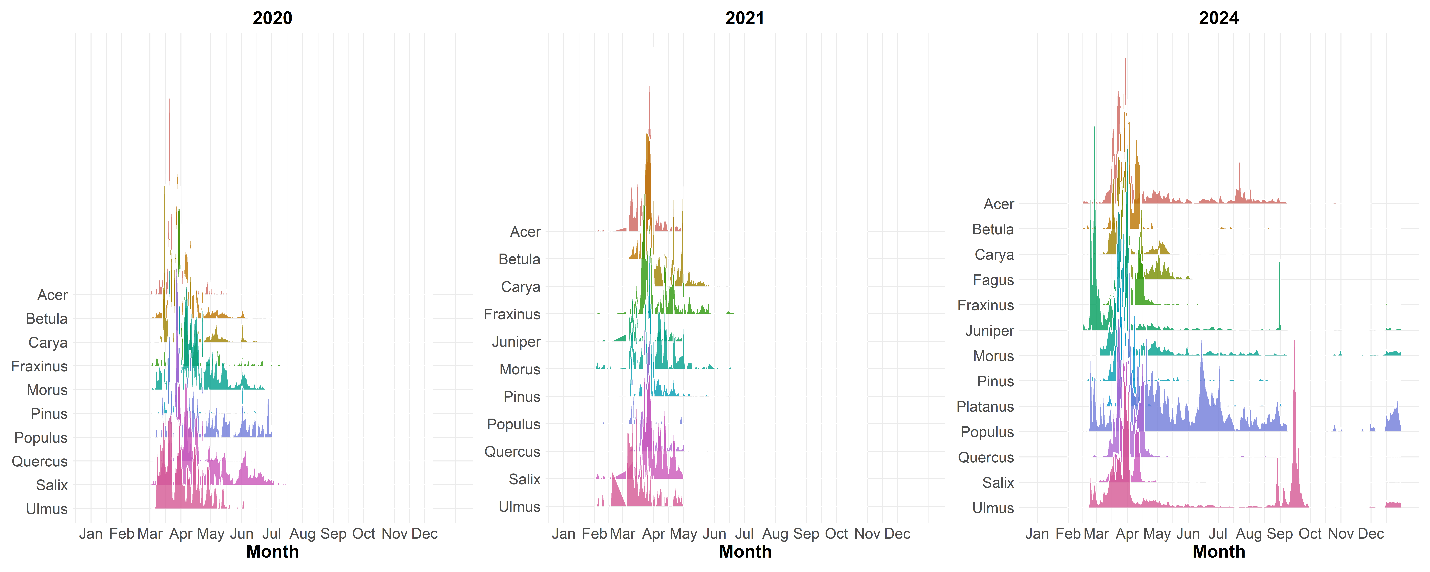

Figure S2. Daily emission factors estimated using logistic growth models by tree taxa and study year.

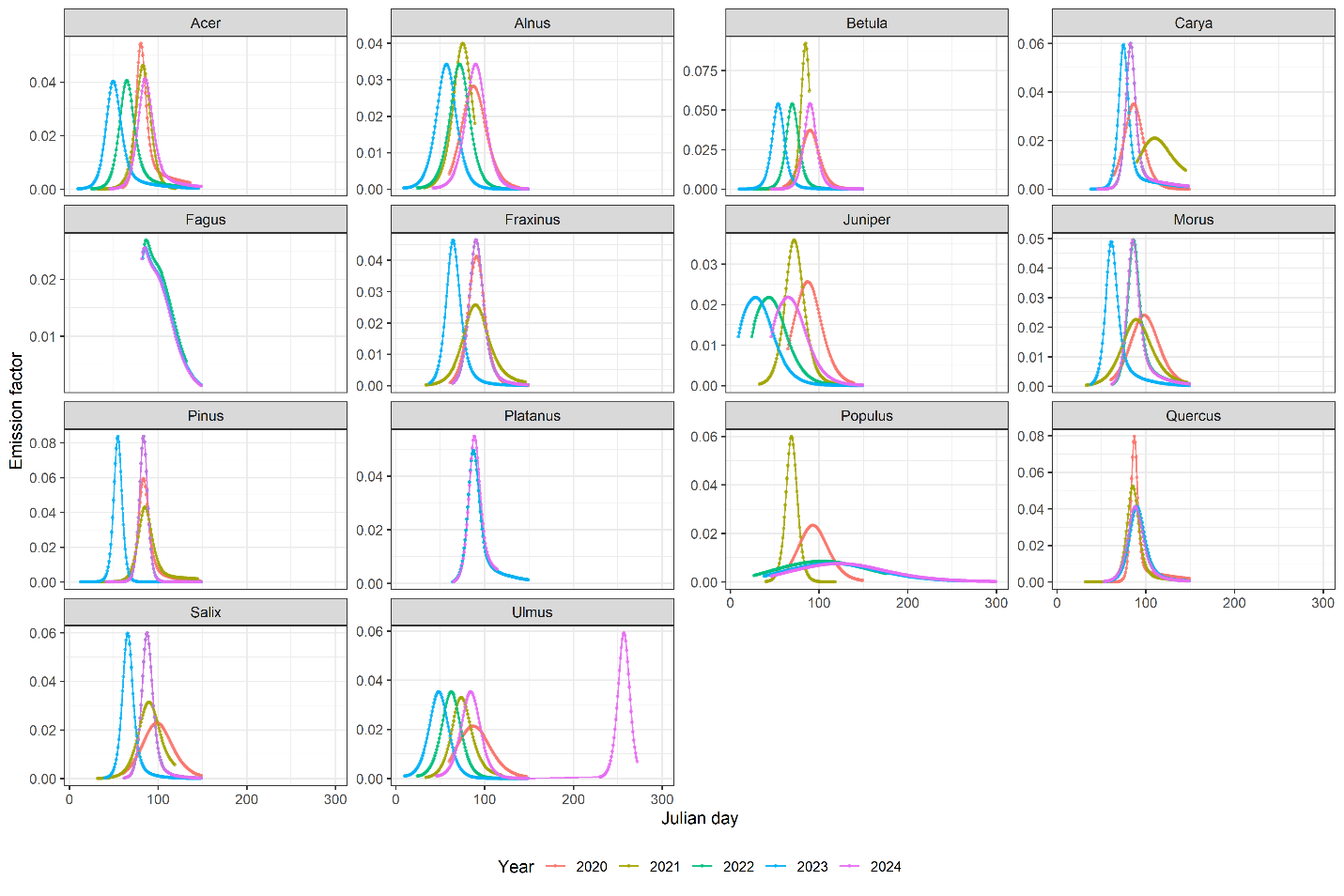

| Table S6: Random 10-fold CV average R^2^ and RMSE/peak for pollen season, 2024. | | | | | | | | | |
| --- | --- | --- | --- | --- | --- | --- | --- | --- | --- |
|  | **RF** | | **XGB** | | **GAM** | | **Temporal R^2^** | | |
| **Tree** | **R^2^** | **RMSE/peak** | **R^2^** | **RMSE/peak** | **R^2^** | **RMSE/peak** | **RF** | **XGB** | **GAM** |
| *Acer* | 0.53 | 0.06 | 0.56 | 0.06 | 0.55 | 0.06 | 0.55 | 0.59 | 0.62 |
| *Betula* | 0.75 | 0.05 | 0.76 | 0.05 | 0.77 | 0.05 | 0.64 | 0.69 | 0.67 |
| *Carya* | 0.7 | 0.05 | 0.66 | 0.05 | 0.67 | 0.05 | 0.67 | 0.55 | 0.54 |
| *Fagus* | 0.25 | 0.06 | 0.33 | 0.05 | 0.29 | 0.06 | 0.47 | 0.53 | 0.52 |
| *Fraxinus* | 0.71 | 0.05 | 0.72 | 0.05 | 0.67 | 0.05 | 0.7 | 0.65 | 0.67 |
| *Juniperus* | 0.44 | 0.05 | 0.63 | 0.04 | 0.67 | 0.04 | 0.41 | 0.4 | 0.43 |
| *Morus* | 0.66 | 0.05 | 0.75 | 0.04 | 0.8 | 0.04 | 0.5 | 0.54 | 0.52 |
| *Pinus* | 0.59 | 0.06 | 0.7 | 0.05 | 0.72 | 0.05 | 0.77 | 0.74 | 0.74 |
| *Platanus* | 0.61 | 0.02 | 0.65 | 0.02 | 0.67 | 0.02 | 0.49 | 0.42 | 0.46 |
| *Populus* | 0.41 | 0.06 | 0.44 | 0.05 | 0.45 | 0.05 | 0.38 | 0.39 | 0.39 |
| *Quercus* | 0.71 | 0.06 | 0.71 | 0.06 | 0.74 | 0.05 | 0.72 | 0.75 | 0.76 |
| *Salix* | 0.70 | 0.06 | 0.73 | 0.05 | 0.71 | 0.05 | 0.6 | 0.57 | 0.5 |
| *Ulmus* (spring only) | 0.78 | 0.02 | 0.93 | 0.01 | 0.94 | 0.01 | 0.52 | 0.70 | 0.69 |
| *Ulmus (two seasons)* | 0.44 | 0.05 | 0.59 | 0.04 | 0.64 | 0.04 | 0.32 | 0.36 | 0.36 |

| Table S7: Random 10-fold CV average R^2^ and RMSE/peak for pollen season, 2021. | | | | | | | | | |
| --- | --- | --- | --- | --- | --- | --- | --- | --- | --- |
|  | **RF** | | **XGB** | | **GAM** | | **Temporal R^2^** | | |
| **Tree** | **R^2^** | **RMSE/peak** | **R^2^** | **RMSE/peak** | **R^2^** | **RMSE/peak** | **RF** | **XGB** | **GAM** |
| *Acer* | 0.48 | 0.14 | 0.56 | 0.12 | 0.51 | 0.12 | 0.49 | 0.53 | 0.56 |
| *Betula* | 0.87 | 0.12 | 0.92 | 0.09 | 0.92 | 0.09 | 0.95 | 0.96 | 0.93 |
| *Carya* | 0.32 | 0.13 | 0.48 | 0.11 | 0.54 | 0.1 | 0.52 | 0.62 | 0.72 |
| *Fraxinus* | 0.54 | 0.09 | 0.61 | 0.08 | 0.56 | 0.08 | 0.43 | 0.57 | 0.5 |
| *Juniperus* | 0.47 | 0.08 | 0.59 | 0.05 | 0.5 | 0.05 | 0.5 | 0.54 | 0.55 |
| *Morus* | 0.48 | 0.09 | 0.44 | 0.1 | 0.49 | 0.1 | 0.39 | 0.37 | 0.47 |
| *Pinus* | 0.75 | 0.1 | 0.81 | 0.08 | 0.83 | 0.08 | 0.67 | 0.72 | 0.73 |
| *Populus* | 0.13 | 0.09 | 0.21 | 0.07 | 0.21 | 0.08 | 0.32 | 0.45 | 0.39 |
| *Quercus* | 0.81 | 0.1 | 0.80 | 0.09 | 0.83 | 0.09 | 0.77 | 0.8 | 0.82 |
| *Salix* | 0.66 | 0.14 | 0.7 | 0.13 | 0.65 | 0.14 | 0.71 | 0.76 | 0.66 |
| *Ulmus* | 0.29 | 0.12 | 0.32 | 0.12 | 0.33 | 0.12 | 0.24 | 0.27 | 0.33 |

| Table S8: Random 10-fold CV R^2^ and RMSE/peak for pollen season, 2020. | | | | | | | | | |
| --- | --- | --- | --- | --- | --- | --- | --- | --- | --- |
|  | **RF** | | **XGB** | | **GAM** | | **Temporal R^2^** | | |
| **Tree** | **R^2^** | **RMSE/peak** | **R^2^** | **RMSE/peak** | **R^2^** | **RMSE/peak** | **RF** | **XGB** | **GAM** |
| *Acer* | 0.63 | 0.04 | 0.48 | 0.05 | 0.49 | 0.06 | 0.58 | 0.49 | 0.50 |
| *Betula* | 0.73 | 0.11 | 0.69 | 0.11 | 0.72 | 0.11 | 0.81 | 0.78 | 0.79 |
| *Carya* | 0.56 | 0.08 | 0.64 | 0.07 | 0.59 | 0.08 | 0.49 | 0.55 | 0.51 |
| *Fraxinus* | 0.69 | 0.09 | 0.67 | 0.09 | 0.58 | 0.10 | 0.62 | 0.62 | 0.57 |
| *Juniperus* | 0.26 | 0.18 | 0.25 | 0.19 | 0.27 | 0.18 | 0.28 | 0.33 | 0.21 |
| *Morus* | 0.74 | 0.10 | 0.77 | 0.09 | 0.77 | 0.09 | 0.50 | 0.55 | 0.58 |
| *Pinus* | 0.53 | 0.09 | 0.51 | 0.09 | 0.50 | 0.10 | 0.59 | 0.48 | 0.51 |
| *Populus* | 0.43 | 0.11 | 0.49 | 0.10 | 0.48 | 0.10 | 0.27 | 0.33 | 0.34 |
| *Quercus* | 0.76 | 0.06 | 0.86 | 0.05 | 0.89 | 0.05 | 0.58 | 0.61 | 0.56 |
| *Salix* | 0.57 | 0.12 | 0.55 | 0.12 | 0.59 | 0.12 | 0.55 | 0.54 | 0.57 |
| *Ulmus* | 0.66 | 0.12 | 0.66 | 0.12 | 0.65 | 0.11 | 0.40 | 0.45 | 0.43 |

| Table S9: Random 10-fold CV average R^2^ and RMSE for pollen count in non-season, 2024. | | | | | | | | | |
| --- | --- | --- | --- | --- | --- | --- | --- | --- | --- |
|  | **RF** | | **XGB** | | **GAM** | | **Temporal R^2^** | | |
| **Tree** | **R^2^** | **RMSE** | **R^2^** | **RMSE** | **R^2^** | **RMSE** | **RF** | **XGB** | **GAM** |
| *Acer* | 0.21 | 6.18 | 0.19 | 6.89 | 0.22 | 7.23 | 0.22 | 0.21 | 0.22 |
| *Betula* | 0.11 | 5.27 | 0.17 | 5.45 | 0.19 | 5.40 | 0.17 | 0.19 | 0.20 |
| *Carya* | 0.12 | 0.52 | 0.10 | 0.51 | 0.14 | 0.59 | 0.21 | 0.25 | 0.23 |
| *Fagus* | 0.29 | 1.49 | 0.47 | 1.52 | 0.37 | 1.55 | 0.74 | 0.77 | 0.73 |
| *Fraxinus* | 0.22 | 2.44 | 0.42 | 2.12 | 0.45 | 2.07 | 0.46 | 0.55 | 0.59 |
| *Juniperus* | 0.04 | 20.21 | 0.07 | 21.89 | 0.10 | 21.64 | 0.16 | 0.15 | 0.23 |
| *Morus* | 0.18 | 22.28 | 0.28 | 20.79 | 0.31 | 20.35 | 0.27 | 0.32 | 0.34 |
| *Pinus* | 0.44 | 16.87 | 0.42 | 18.47 | 0.40 | 18.90 | 0.26 | 0.27 | 0.24 |
| *Platanus* | 0.56 | 439.48 | 0.63 | 393.25 | 0.66 | 364.89 | 0.53 | 0.61 | 0.64 |
| *Populus* | 0.23 | 2.32 | 0.33 | 2.15 | 0.37 | 1.94 | 0.34 | 0.46 | 0.43 |
| *Quercus* | 0.24 | 9.10 | 0.23 | 8.71 | 0.21 | 9.40 | 0.26 | 0.23 | 0.25 |
| *Salix* | 0.17 | 4.15 | 0.25 | 3.93 | 0.36 | 3.48 | 0.29 | 0.31 | 0.38 |
| *Ulmus* | 0.49 | 6.57 | 0.63 | 5.52 | 0.60 | 5.55 | 0.26 | 0.35 | 0.32 |

| Table S10: Random 10-fold CV R^2^ and RMSE/peak for pollen count in non-season, 2021. For some taxa, during random 10-fold cross-validation, the testing sets contained fewer than five days with non-zero pollen counts; as a result, the linear regression used to estimate R² did not converge. | | | | | | | | | |
| --- | --- | --- | --- | --- | --- | --- | --- | --- | --- |
|  | RF | | XGB | | **GAM** | | **Temporal R^2^** | | |
| **Tree** | **R^2^** | **RMSE/peak** | **R^2^** | **RMSE/peak** | **R^2^** | **RMSE/peak** | **RF** | **XGB** | **GAM** |
| *Acer* | Few non-zero count | | | | | | | | |
| *Betula* | 0.97 | 11.19 | 0.97 | 16.54 | 1.00 | 17.54 | 0.97 | 0.97 | 1.00 |
| *Carya* | 0.66 | 1.43 | 0.59 | 1.61 | 0.67 | 2.85 | 0.97 | 0.87 | 1.00 |
| *Fraxinus* | Few non-zero count | | | | | | | | |
| *Juniperus* | 0.63 | 7.78 | 0.86 | 11.10 | 0.90 | 6.90 | 0.82 | 0.84 | 0.92 |
| *Morus* | Few non-zero count | | | | | | | | |
| *Pinus* | Few non-zero count | | | | | | | | |
| *Populus* | Few non-zero count | | | | | | | | |
| *Quercus* | Few non-zero count | | | | | | | | |
| *Salix* | 0.50 | 3.12 | 0.91 | 5.94 | 0.94 | 6.81 | 0.85 | 0.94 | 0.93 |
| *Ulmus* | Few non-zero count | | | | | | | | |

| Table S11: Random 10-fold CV average R^2^ and RMSE/peak for pollen count in non-season, 2024. | | | | | | | | | |
| --- | --- | --- | --- | --- | --- | --- | --- | --- | --- |
|  | **RF** | | **XGB** | | **GAM** | | **Temporal R^2^** | | |
| **Tree** | **R^2^** | **RMSE** | **R^2^** | **RMSE** | **R^2^** | **RMSE** | **RF** | **XGB** | **GAM** |
| *Acer* | 0.07 | 0.33 | 0.01 | 0.35 | 0.05 | 0.30 | 0.42 | 0.03 | 0.05 |
| *Betula* | 0.41 | 7.55 | 0.57 | 5.93 | 0.53 | 6.21 | 0.48 | 0.57 | 0.53 |
| *Carya* | 0.20 | 17.14 | 0.19 | 17.35 | 0.21 | 16.60 | 0.46 | 0.37 | 0.21 |
| *Fraxinus* | 0.16 | 1.30 | 0.22 | 1.34 | 0.23 | 1.33 | 0.30 | 0.34 | 0.23 |
| *Juniperus* | Few non-zero count | | | | | | | | |
| *Morus* | 0.51 | 3.48 | 0.54 | 3.44 | 0.40 | 4.72 | 0.50 | 0.53 | 0.40 |
| *Pinus* | 0.31 | 14.44 | 0.22 | 14.83 | 0.18 | 15.82 | 0.37 | 0.24 | 0.18 |
| *Populus* | 0.00 | 0.20 | 0.00 | 0.16 | 0.00 | 0.30 | 0.95 | 0.01 | 0.00 |
| *Quercus* | 0.06 | 0.47 | 0.05 | 0.44 | 0.33 | 0.40 | 0.28 | 0.35 | 0.33 |
| *Salix* | 0.58 | 8.26 | 0.74 | 6.20 | 0.77 | 5.89 | 0.58 | 0.64 | 0.77 |
| *Ulmus* | 0.31 | 0.44 | 0.23 | 0.47 | 0.13 | 0.32 | 0.32 | 0.23 | 0.13 |

| Table S12. Random 10-fold CV R^2^ and RMSE for cross-year predictions and compared with same-year emission factors, 2021. | | | | | | | | | | | | |
| --- | --- | --- | --- | --- | --- | --- | --- | --- | --- | --- | --- | --- |
|  | **RF** | | | | **XGB** | | | | **GAM** | | | |
| **Tree** | **R^2^** | **Diff.** | **RMSE** | **Diff.** | **R^2^** | **Diff.** | **RMSE** | **Diff.** | **R^2^** | **Diff.** | **RMSE** | **Diff.** |
| *Acer* | 0.59 | +0.11 | 40.49 | -9.35 | 0.61 | +0.05 | 37.14 | -6.30 | 0.66 | +0.16 | 36.49 | -7.29 |
| *Betula* | 0.76 | -0.11 | 142.52 | -27.37 | 0.90 | -0.02 | 102.92 | -36.24 | 0.89 | -0.03 | 106.47 | -29.47 |
| *Carya* | 0.38 | +0.06 | 55.15 | -14.49 | 0.41 | -0.07 | 60.50 | -2.32 | 0.53 | -0.02 | 53.42 | +0.90 |
| *Fraxinus* | 0.64 | +0.11 | 45.95 | -7.33 | 0.61 | -0.01 | 48.85 | +2.28 | 0.65 | +0.09 | 49.30 | +1.67 |
| *Juniperus* | 0.39 | -0.08 | 350.12 | -50.98 | 0.48 | -0.10 | 311.74 | +74.81 | 0.47 | -0.03 | 285.42 | +24.17 |
| *Morus* | 0.51 | +0.03 | 31.43 | +1.93 | 0.45 | +0.01 | 33.99 | +2.14 | 0.40 | -0.09 | 34.28 | +3.50 |
| *Pinus* | 0.54 | -0.21 | 723.53 | +65.14 | 0.73 | -0.08 | 521.78 | -4.21 | 0.72 | -0.11 | 498.66 | -3.55 |
| *Populus* | 0.06 | -0.07 | 10.35 | -3.99 | 0.18 | -0.02 | 11.23 | -0.15 | 0.17 | -0.04 | 10.48 | -1.84 |
| *Quercus* | 0.88 | +0.08 | 866.09 | -46.25 | 0.93 | +0.13 | 724.55 | -159.18 | 0.86 | +0.03 | 840.96 | -56.59 |
| *Salix* | 0.67 | +0.01 | 82.73 | -4.52 | 0.70 | +0.00 | 81.83 | +2.41 | 0.69 | +0.04 | 75.80 | -9.27 |
| *Ulmus* | 0.34 | +0.05 | 12.80 | -2.66 | 0.43 | +0.11 | 11.51 | -3.11 | 0.45 | +0.12 | 11.70 | -3.96 |

Figure S3. Time series plots of XGBoost-predicted pollen count among the testing sets from 10-fold random CV approach for the pollen season in 2020 and 2021.

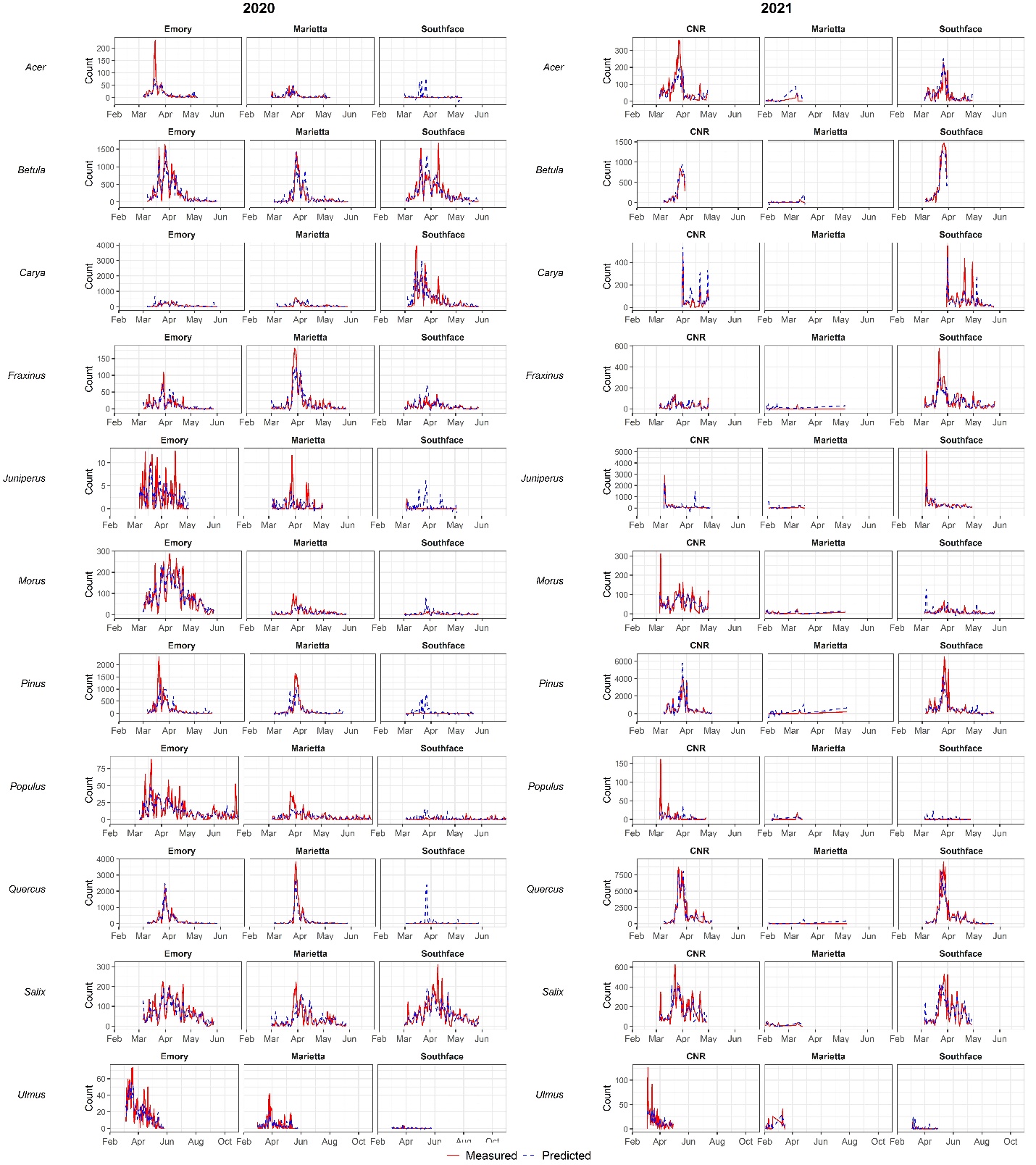

Figure S4. Time series plots of XGBoost-predicted pollen count among the testing sets from 10-fold random CV approach for the pollen season in 2024.

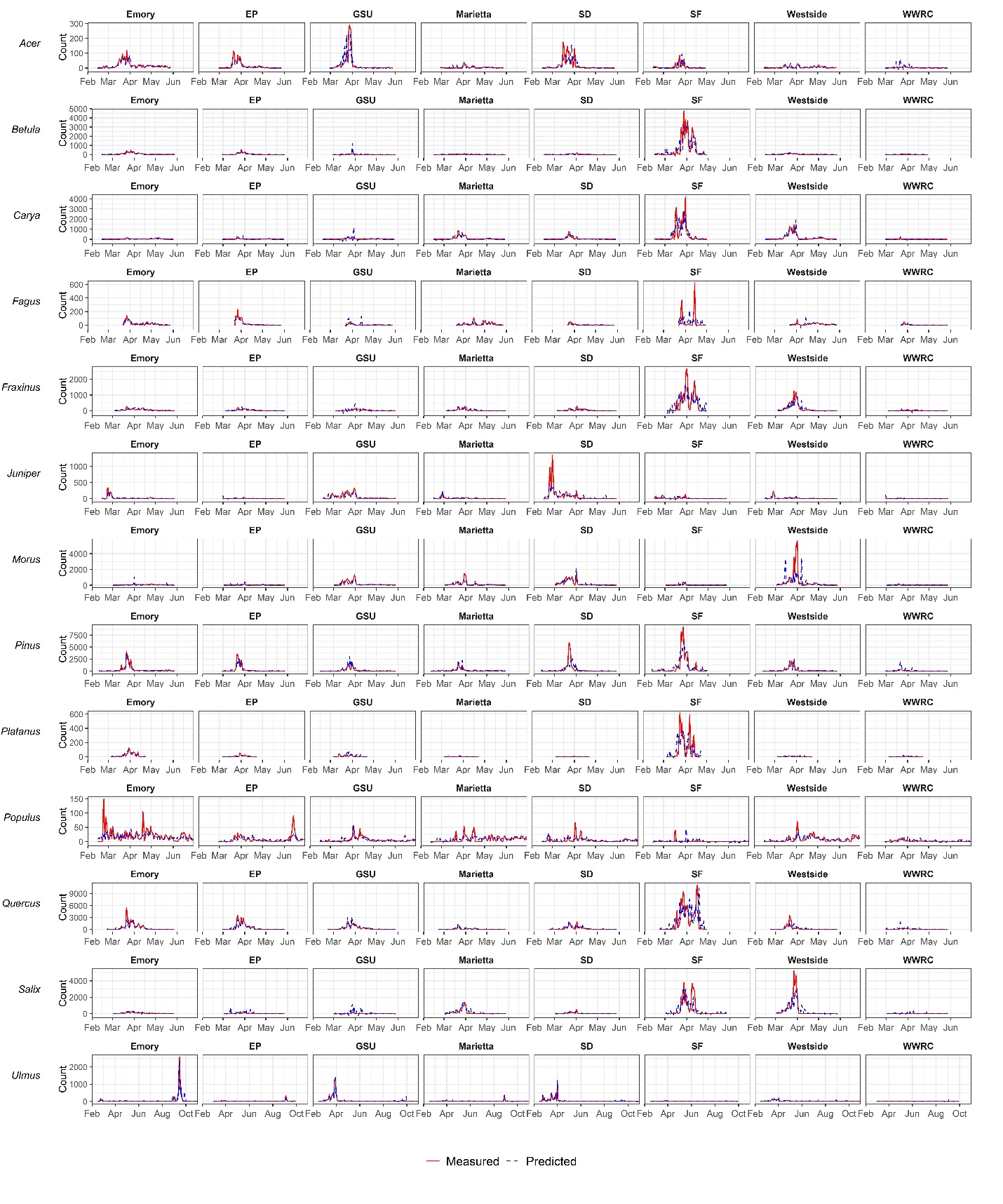

Gregory, Philip Herries. *The microbiology of the atmosphere*. London: Leonard Hill, 1961.
